# Multiscale Complexity as a Basis for Functional Brain Network Construction

**DOI:** 10.64898/2026.03.28.715014

**Authors:** Amir Hossein Ghaderi, Mary Helen Immordino-Yang

## Abstract

Functional brain networks are conventionally constructed using measures of direct temporal synchrony between neural signals, implicitly restricting connectivity to scale-specific interactions. Here, we introduce an alternative framework in which interregional similarity is defined through correlations between multiscale entropy (MSE) profiles, enabling network construction based on scale-dependent dynamical structure rather than instantaneous alignment. Using resting-state fMRI data from the Human Connectome Project (N = 1003), we systematically compare MSE-based networks with conventional time-series–based networks across conventional/spectral graph-theoretical, and information-theoretic measures. We show that MSE-based networks exhibit stronger modular organization, enhanced local segregation, and distinct global integration patterns, reflecting a reorganization of functional architecture when multiscale dynamics are taken into account. Importantly, MSE-based networks demonstrate substantially greater sensitivity to biologically meaningful variability, revealing robust and reproducible sex differences across multiple network measures, in contrast to the limited and inconsistent effects observed in conventional networks. These findings suggest that multiscale representations provide a more informative and biologically grounded basis for functional brain network construction, capturing aspects of neural organization that are not accessible through direct synchrony alone.

## I. INTRODUCTION

Functional brain networks are typically inferred by assigning edges according to measures of pairwise synchrony between regional neural signals[1, 2]. Such measures, whether linear or nonlinear, quantify statistical dependence through direct temporal or phase alignment and therefore encode functional coupling as a form of co-ordinated activity at specific time or frequency scales[3, 4]. This construction implicitly assumes that interregional interactions are sufficiently characterized by synchronization within a limited dynamical bandwidth[2, 4]. However, neural activity is intrinsically multiscale, exhibiting structured fluctuations across a broad hierarchy of temporal and frequency[5, 6]. Rather than operating within a single dominant scale, the brain engages in concurrent dynamics spanning slow and fast processes, with interactions distributed across multiple frequency bands and time scales[7, 8]. In this context, direct synchrony measures act as scale-selective operators, privileging temporal coincidence within predefined bands and potentially overlooking similarities that emerge only when the full multiscale organization of the signals is considered.

From a dynamical systems standpoint, two neural processes may display weak or negligible pairwise synchrony while nevertheless sharing comparable scale-dependent statistical structure, such as similar entropy decay, complexity profiles, or hierarchical variability across temporal resolutions[8]. Consequently, networks derived from direct synchrony estimates may conflate local coordination at specific scales with global similarity in dynamical organization, thereby providing an incomplete representation of multiscale neural dynamics[9, 10]. Defining interregional similarity instead through correlations between multiscale descriptors, such as multiscale entropy (MSE), offers a formal alternative in which network edges encode similarities in scale-dependent structure rather than instantaneous alignment. This formulation reframes functional brain networks as graphs of dynamical equivalence across scales, aligning more naturally with the view of the brain as a frequency-multiplexed, hierarchically organized complex system[5, 8, 11].

MSE analysis has been widely employed to characterize the intrinsic complexity of neural signals by quantifying their irregularity across progressively coarser temporal scales[8, 11, 12]. In the context of brain dynamics, MSE has consistently revealed that neural activity cannot be adequately described by single-scale measures of variability or predictability. Instead, healthy and adaptive brain function is associated with structured entropy profiles spanning both fine and coarse temporal scales, reflecting the coexistence of fast, locally fluctuating processes and slower, integrative dynamics. Fine-scale entropy is commonly interpreted as indexing short-range, high-frequency fluctuations that are thought to reflect local neural information processing. In contrast, coarse-scale entropy captures long-range temporal dependencies that are integrated across distributed neural populations[8]. Importantly, alterations in these scale-dependent entropy profiles have been linked to changes in cognitive state, development, and pathology, suggesting that MSE encodes functionally meaningful aspects of neural organization rather than incidental signal variability[8, 13, 14]. From a network perspective, these scale-specific entropy characteristics imply that functional interactions between brain regions[5] may be more naturally expressed in terms of similarities in their MSE trajectories than through direct temporal synchronization. By correlating entropy profiles across regions, adjacency matrices can be constructed in which edge weights reflect similarity in scale-dependent complexity, thereby embedding multiscale and nonlinear features of neural dynamics directly into network architecture.

In the present work, we adopt this multiscale perspective to construct functional brain networks from empirical fMRI data obtained from the human connectome project (HCP-1200)[15]. For each subject, functional networks are independently derived using two complementary formulations: a conventional time-series–based (TS-based) approach, in which adjacency weights encode direct statistical dependence between regional signals, and a MSE–based approach, in which inter-regional similarity is defined through correlations between scale-dependent entropy profiles. Using randomized null models, network measures are normalized to control for trivial effects of network density and weight distribution, allowing a principled comparison between the two constructions[16].

Based on the premise that MSE–based networks explicitly encode scale-dependent and nonlinear properties of neural dynamics[8], we hypothesize that such networks are better suited to highlight mesoscale organization and biologically meaningful variability in brain function than TS–based networks. Furthermore, we posit that graph-theoretical measures, including both conventional and spectral descriptors, as well as network-level Shannon entropy, will more effectively differentiate functional brain organization across groups when applied to MSE–based networks, compared to TS–based networks. As a well-defined reference for group-level comparison, we consider sex as a basic biological feature to evaluate the sensitivity of different network constructions to structured variability in brain organization. The following sections describe the construction of these networks, and the analytical framework used to evaluate these hypotheses.

## II. DATASET, FMRI PREPROCESSING, AND NODES DEFINITION

We analyzed resting-state fMRI data from the HCP-1200 release of the Human Connectome Project (HCP)[15], comprising high-resolution neuroimaging data acquired under harmonized protocols in a large cohort of healthy young adults[17]. The fMRI sample used in the present study consisted of N= 1003 participants (534 females)[17]. Analyses were performed on HCP-distributed minimally preprocessed data, which included spatial distortion correction, motion correction, and cross-subject normalization. Structured artifacts were removed using an independent component analysis (ICA)–based denoising procedure[18, 19].

Functional network nodes were defined via group-ICA decomposition with dimensionality *D* = 300, corresponding to the maximum number of components available in the HCP extensively processed fMRI dataset. This high-dimensional representation was chosen to maximize network size and spatial resolution, thereby enabling a more detailed characterization of large-scale functional organization. Each ICA component represents a spatially independent functional pattern spanning cortical and sub-cortical gray-matter regions and was treated as a node in the functional network. For each subject, a representative BOLD time series was extracted for every component via dual regression, yielding a multivariate signal 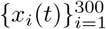 describing region-specific dynamics. This ICA-derived node set was held fixed across all analyses.

## III. MSE ESTIMATION

To quantify the scale-dependent complexity of regional brain activity, MSE was computed independently for the BOLD time series associated with each network node. Let *x*_*i*_(*t*) denote the BOLD time series of node *i*, with *t* = 1, …, *T*. For each node *i* and for a given scale factor *τ*, a coarse-grained time series 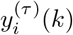 was constructed as[20]

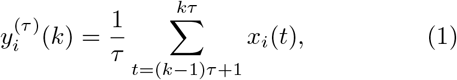

where *k* = 1, …, ⌊*T/τ*⌋. This coarse-graining procedure was applied separately to the time series of each node, thereby isolating temporal dynamics at progressively coarser scales for each region.

For every coarse-grained time series 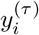, we computed the sample entropy SampEn(*m, r*), which estimates the negative logarithm of the conditional probability that two sequences of length *m* that are similar within tolerance *r* remain similar when extended to length *m* + 1. Sample entropy is defined as[20]

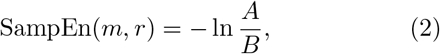

where *B* denotes the number of pairs of embedded vectors of dimension *m* whose distance is less than *r*, and *A* is the corresponding number for dimension *m* + 1. Distances were calculated using the maximum norm and tol-erance *r* was defined as a fixed proportion of the standard deviation of the original node-specific time series.

**FIG. 1.**
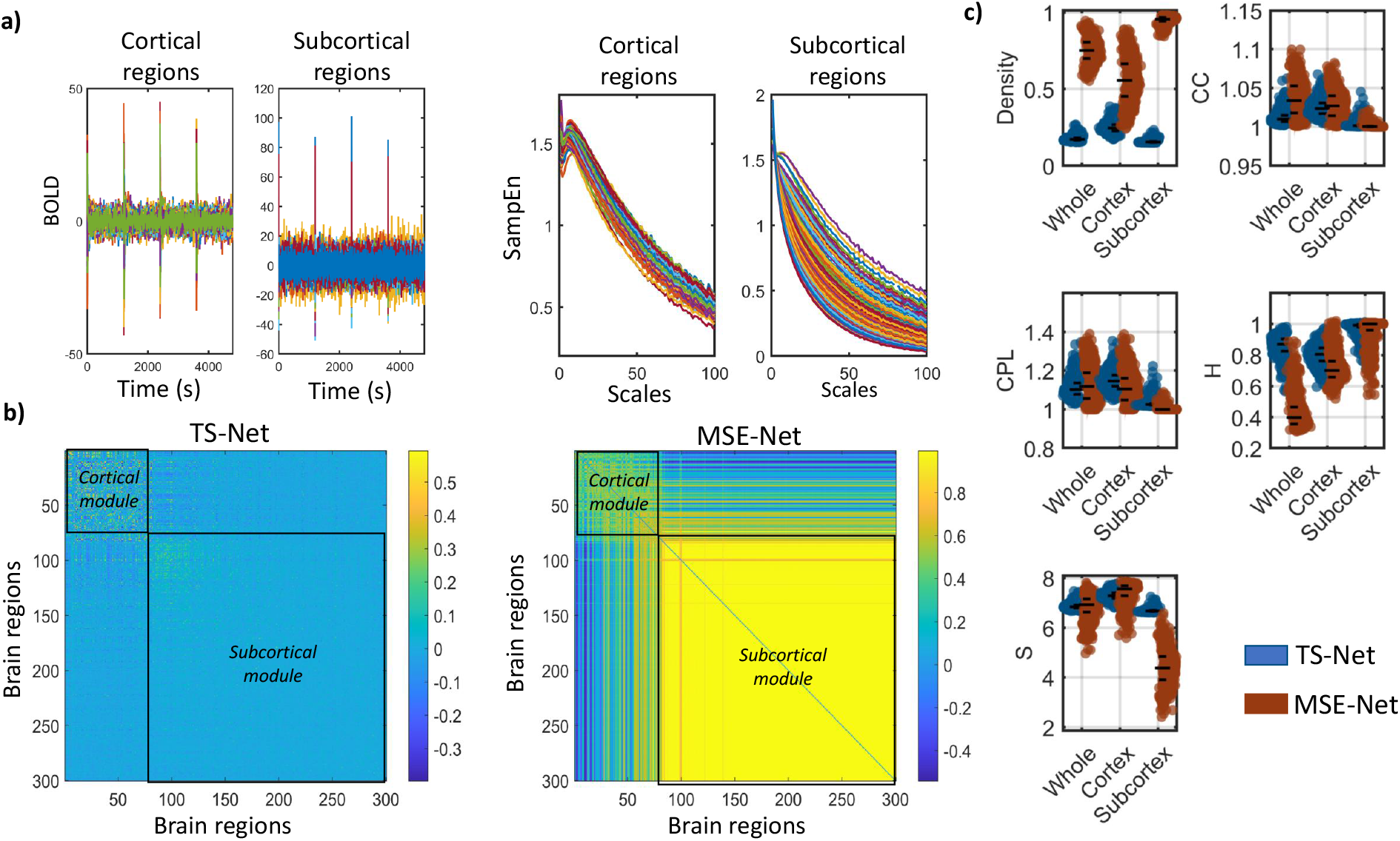
Schematic illustration of functional brain network construction using time-series–based (TS) and multiscale entropy–based (MSE) formulations. (a) In TS-Nets, edges are defined by the absolute Pearson correlation between regional BOLD time series. In MSE-Nets, each region is first represented by its multiscale entropy (MSE) profile across scales (1 to 100), and edges are defined by the similarity (absolute Pearson correlation) between these profiles. (b) Resulting network representations reveal differences in mesoscale organization, with both approaches yielding modular structures broadly corresponding to cortical and subcortical divisions, but with stronger modular separation in MSE-based networks. (c)Quantitative comparison of graph-theoretical measures between TS-Nets and MSE-Nets for whole-brain, cortical, and subcortical systems, highlighting systematic differences in connection density (*CD*), clustering coefficient (*CC*), characteristic path length (*CPL*), graph energy (*H*), and entropy (*S*) which are associated with connectivity strength, segregation, integration, synchronization stability, and edge-weight heterogeneity, respectively.

Repeating this procedure across a range of scale factors *τ* = 1, …, *τ*_max_ yields, for each node *i*, a multiscale entropy profile[20]

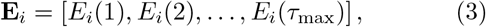

where *E*_*i*_(*τ*) denotes the sample entropy of node *i* at scale *τ*. Each node is therefore represented by a scale-dependent entropy vector encoding the hierarchical temporal organization of its activity.

We selected *m* and *r* parameters based on previous studies and our data-driven approach, respectively. According to many previous fMRI studies, MSE was computed using sample entropy with embedding dimension fixed to *m* = 2[5, 21, 22]. The tolerance parameter *r* was selected using a data-driven validity criterion motivated by the mathematical properties of SampEn(*m, r*) estimation. For small values of *r*, the tolerance region becomes overly restrictive, such that for some nodes no pairs of embedded vectors satisfy the similarity criterion.

In these cases, the conditional probabilities required for SampEn(*m, r*) estimation cannot be evaluated, resulting in undefined entropy values for those nodes.

To account for this limitation, we systematically increased *r* starting from *r* = 0.1 and, for each candidate value, quantified the proportion of nodes for which sample entropy could not be computed across the resulting entropy matrices. We defined an acceptable tolerance as one for which fewer than 5% of nodes yielded undefined entropy values, indicating that a sufficient number of template matches existed for reliable estimation. Based on this criterion, the smallest admissible tolerance was identified as *r* = 0.27, which balanced computational stability against excessive smoothing of the underlying dynamics.

## IV. ADJACENCY MATRICES AND MODULAR ORGANIZATION

Functional brain networks were constructed using two alternative and explicitly comparative definitions of interregional similarity. In the first formulation, adjacency matrices were derived from absolute values of Pearson correlations between node-specific BOLD time series, yielding TS–Nets 1:

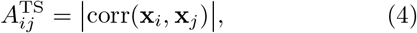

where **x**_*i*_ and **x**_*j*_ are the BOLD time series of nodes *i* and *j*, respectively.

In the second formulation, adjacency matrices were constructed by computing similarity between MSE profiles (absolute values of Pearson correlations across scales 1–100) associated with each node, resulting in MSE–Nets (1):

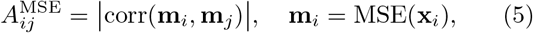

where **m**_*i*_ denotes MSE profile of node *i*. The node set was held fixed across both formulations and defined by the high-dimensional group-ICA parcellation, ensuring that observed differences in network organization arise solely from the definition of edge weights.

To characterize mesoscale organization in each network representation, we employed a modularity optimization framework[23, 24]. Modularity quantifies the extent to which a network can be partitioned into modules with stronger within-module connectivity than expected under an appropriate null model. The modularity quality function is defined as[24]

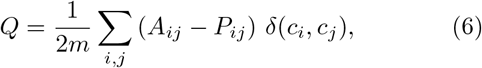

where *A*_*ij*_ denotes the observed adjacency matrix, *P*_*ij*_ represents the expected connectivity under a null model with preserved node strengths, *m* is the total edge weight in the network, *c*_*i*_ is the module assignment of node *i*, and *δ*(*c*_*i*_, *c*_*j*_) equals unity when nodes *i* and *j* belong to the same module and zero otherwise. Maximization of *Q* yields a partition that emphasizes nontrivial community structure beyond global connectivity effects. We performed this analysis separately on the averaged TS-Nets and averaged MSE-Nets. As shown in 2, for TS-Nets, the maximum *Q* allowing at least two separate modules was 0.078, whereas for MSE-Nsts, this value was 0.096. Further investigation revealed that these two modules predominantly corresponded to largely separate cortical and subcortical regions in both TS-Nets and MSE-Nets. The higher modularity ratio observed in MSE–Nets indicates that MSE-based connectivity more strongly captures the cortical–subcortical modular structure of the brain compared to direct temporal synchrony.

**FIG. 2.**
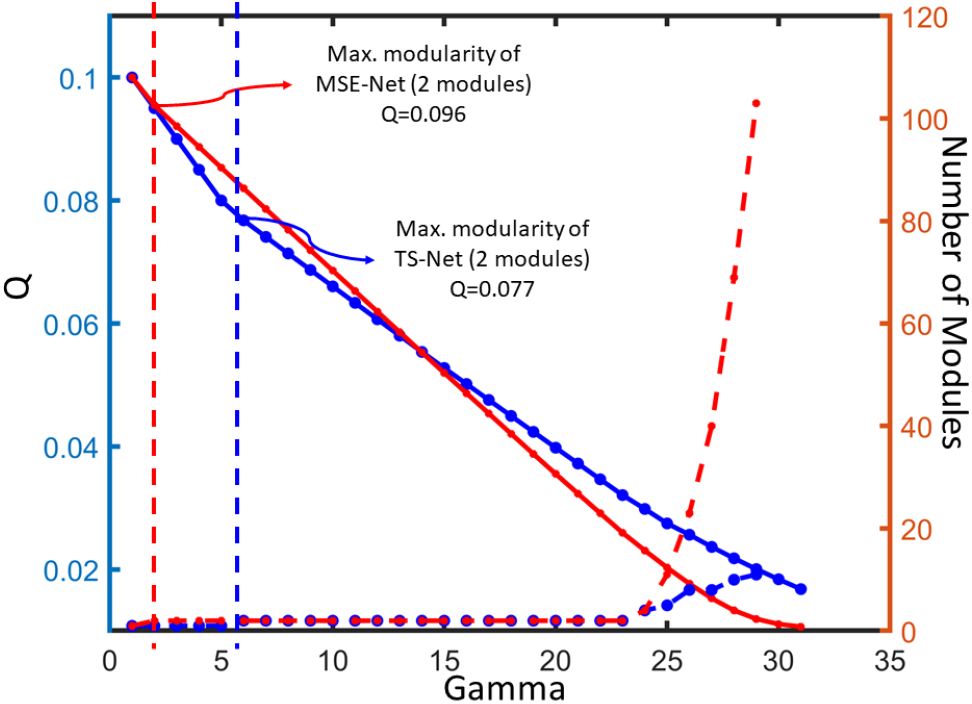
Modular organization of TS-based and MSE-based functional brain networks across resolution scales (Gamma). Networks were constructed from averaged adjacency matrices and partitioned using modularity maximization while varying the resolution parameter (*γ*), which controls the granularity of community detection. For each network type, we identified the maximum modularity value (*Q*) at which the network partitions into at least two distinct modules. This value was lower for TS-Nets (*Q* = 0.078) than for MSE-Nets (*Q* = 0.096), indicating stronger community structure in MSE-based networks. In both cases, the resulting modules predominantly correspond to a separation between cortical and subcortical regions.

## V. GRAPH-THEORETICAL MEASURES AND NETWORK DIFFERENCES

To compare TS–Nets and MSE–Nets properties, we computed five weighted graph measures capturing complementary aspects of network organization: connection density (*CD*), clustering coefficient (*CC*), characteristic path length (*CPL*), graph energy (*H*), and network-level Shannon entropy (*S*). All graphs were modeled as weighted and undirected, and absolute edge weights were used to ensure non-negativity.

To control for trivial effects arising from weight distribution and network density, we employed a null-model normalization[16]. Specifically, for each network, we generated randomized counterparts by shuffling edge weights while preserving the overall weight distribution. *CC, CPL*, and *H* were computed for these shuffled networks, and the corresponding values obtained from the empirical networks were normalized relative to their null-model counterparts. This procedure ensures that potential differences in the average edge weights across networks do not unduly influence the graph-theoretical measures, thereby isolating nontrivial topological and spectral effects from simple weight magnitude differences. *CD* and *S* were not subjected to null-model normalization. Because weight shuffling preserves the global weight distribution, both *CD* and *S* remain invariant under this randomization procedure and therefore cannot be mean-ingfully normalized in this framework. For Shannon entropy, edge weights were instead normalized with respect to their maximum value prior to probability estimation.

**FIG. 3.**
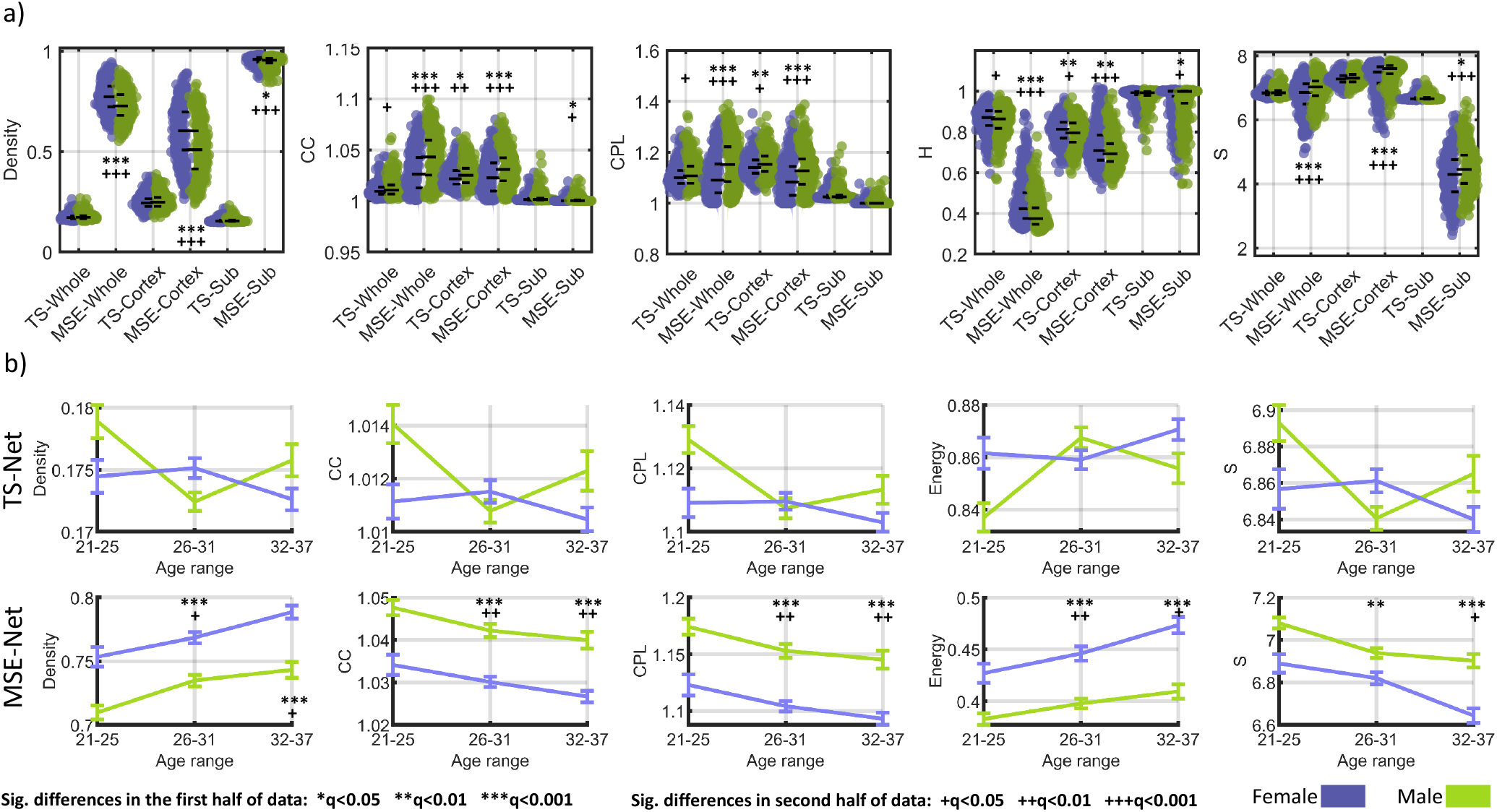
Sex differences in graph-theoretical measures for TS-based and MSE-based functional brain networks. Significant differences are indicated for the first half (*, **, ***) and second half (+, ++, +++) of the data, corresponding to increasing levels of statistical significance. (a) Group comparisons between females and males for connection density (*CD*), clustering coefficient (*CC*), characteristic path length (*CPL*), graph energy (*H*), and Shannon entropy (*S*), computed for whole-brain, cortical, and subcortical networks. (b) Age-stratified analysis of sex differences across three cohorts (21–25, 26–31, and 32–37 years), demonstrating the consistency and robustness of effects across different age ranges. MSE-based networks show robust and consistent sex differences across measures, network scales, and age cohorts, whereas TS-based networks exhibit limited and less reproducible effects.

Statistical comparisons between groups and network types were performed using nonparametric permutation-based *t*-tests (with 5000 random shuffels)[25]. To account for multiple comparisons across graph measures, *p*-values were corrected using the false discovery rate (FDR) procedure.

1 shows the results of the comparison between TS–Nets and MSE–Nets for the whole-brain network as well as for cortical and subcortical subnetworks.

### a. Connection density (CD)

For weighted networks, connection density was defined as the average edge weight across all node pairs[2]:

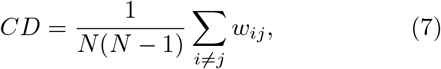

where *N* denotes the number of nodes and *w*_*ij*_ represents the (absolute) weight of the edge between nodes *i* and *j*.

Across all three network definitions (whole-brain, cortical, and subcortical), MSE–Nets exhibited higher *CD* than TS–Nets, indicating systematically stronger average connectivity under the MSE-based similarity construction. This finding indicates that interregional similarity is stronger when defined in terms of MSE profiles than when defined via direct temporal correlations.

### b. Clustering coefficient (CC)

The weighted clustering coefficient of node *i* was defined as[26]

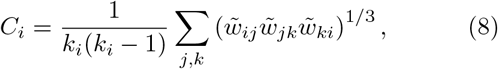

where *k*_*i*_ is the degree of node *i*, and 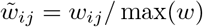 denotes normalized edge weights. The global clustering coefficient was obtained by averaging over all nodes:

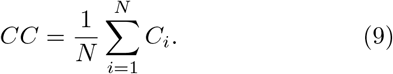

Same as *CD*, the *CC* was higher in MSE–Nets for the whole-brain network and for both cortical and subcortical subnetworks. Given that CC indexes local segregation[2], this result indicates stronger local clustering in MSE-based networks.

**TABLE I.** Statistical comparison of graph-theoretical measures between TS-based (TS-Nets) and MSE-based (MSE-Nets) functional brain networks. Values are reported as t-statistics and corresponding FDR-corrected p-values (t/q) obtained from permutation-based t-tests. Results are shown for whole-brain, cortical, and subcortical networks across two independent analyses (random splits of the dataset). Graph measures include connection density (*CD*), clustering coefficient (*CC*), characteristic path length (*CPL*), graph energy (*H*), and Shannon entropy (*S*). Significant differences (*q <* 0.05) are indicated in **bold**.

|  |  | TS-Whole (t/q) | MSE-Whole (t/q) | TS-Cortex (t/q) | MSE-Cortex (t/q) | TS-Sub (t/q) | MSE-Sub (t/q) |
| --- | --- | --- | --- | --- | --- | --- | --- |
| Density | Analysis 1 | -0.46/.714 | **6.86/ < .001** | -1.28/.232 | **7.45/ < .001** | 0.10/.944 | **2.54/.025** |
|  | Analysis 2 | -1.83/.096 | **7.26/ < .001** | -1.94/.077 | **7.91/ < .001** | -1.51/.167 | **4.20/ < .001** |
| CC | Analysis 1 | -1.21/.259 | **-7.21/ < .001** | **-3.22/.004** | **-4.19/ < .001** | -0.07/.945 | **-2.24/.040** |
|  | Analysis 2 | **-2.49/.018** | **-7.41/ < .001** | **-2.65/.015** | **-5.51/ < .001** | -1.96/.077 | **-2.38/.027** |
| CPL | Analysis 1 | -1.88/.085 | **-7.34/ < .001** | **-3.71/.001** | **-5.09/ < .001** | -0.12/.944 | -1.87/.090 |
|  | Analysis 2 | **-2.44/.027** | **-7.44/ < .001** | **-2.59/.022** | **-5.79/ < .001** | -1.77/.104 | -2.05/.060 |
| H | Analysis 1 | 1.26/.241 | **6.83/ < .001** | **3.13/.003** | **3.12/.003** | -0.10/.944 | **2.32/.035** |
|  | Analysis 2 | **2.15/.042** | **7.13/ < .001** | **2.48/.025** | **3.57/ < .001** | 1.69/.113 | **2.83/.013** |
| S | Analysis 1 | -0.67/.554 | **-4.85/ < .001** | -1.76/.107 | **-5.25/ < .001** | 0.09/.944 | **-2.46/.028** |
|  | Analysis 2 | -1.82/.095 | **-6.44/ < .001** | -1.84/.090 | **-5.71/ < .001** | -1.50/.162 | **-4.67/ < .001** |

**TABLE II.** Statistical comparison of graph-theoretical measures between TS-based (TS-Nets) and MSE-based (MSE-Nets) functional brain networks within whole-brain across different age-ranges. Values are reported as *t*-statistics and corresponding FDR-corrected *p*-values (t/q) obtained from permutation-based *t*-tests. Results are presented for cortical and subcortical subnetworks in three age cohorts (21 −25, 26 −31, and 32 −37 years) across two independent dataset splits (Analysis 1 and Analysis 2). Graph measures include connection density (CD), clustering coefficient (CC), characteristic path length (CPL), Shannon entropy (S), and graph energy (H). Significant differences (*q <* 0.05) are indicated in **bold**.

| Age-range |  | Density (t/q) | CC (t/q) | CPL (t/q) | H (t/q) | S (t/q) |
| --- | --- | --- | --- | --- | --- | --- |
| **TS-Whole** |  |  |  |  |  |  |
| 21–25 | Analysis 1 | -0.01/.993 | -0.50/.768 | -0.64/.720 | 0.41/.789 | -0.04/.993 |
|  | Analysis 2 | -2.04/.122 | -2.16/.122 | -2.45/.122 | 2.33/.122 | -2.14/.122 |
| 26–31 | Analysis 1 | 2.21/.279 | 1.44/.411 | 0.82/.623 | -1.51/.411 | 2.04/.279 |
|  | Analysis 2 | 1.34/.423 | 0.26/.859 | -0.01/.994 | -0.79/.531 | 1.31/.423 |
| 32–37 | Analysis 1 | -1.09/.460 | -1.24/.411 | -1.26/.411 | 1.30/.411 | -1.21/.411 |
|  | Analysis 2 | -1.17/.429 | -1.13/.429 | -0.74/.532 | 1.00/.438 | -1.06/.429 |
| **MSE-Whole** |  |  |  |  |  |  |
| 21–25 | Analysis 1 | 1.27/.227 | -1.44/.201 | -1.36/.212 | 1.42/.201 | -0.60/.548 |
|  | Analysis 2 | **3.75/.002** | **-3.32/.004** | **-3.33/.004** | **3.89/ < .001** | **-3.32/.004** |
| 26–31 | Analysis 1 | **4.94/ < .001** | **-5.40/ < .001** | **-5.56/ < .001** | **4.95/ < .001** | **-3.03/.004** |
|  | Analysis 2 | **2.57/.016** | **-3.34/.004** | **-3.47/.002** | **3.07/.005** | -1.40/.156 |
| 32–37 | Analysis 1 | **3.89/ < .001** | **-3.95/ < .001** | **-3.96/ < .001** | **4.28/ < .001** | **-3.51/.001** |
|  | Analysis 2 | **2.65/.011** | **-3.05/.005** | **-3.00/.005** | **2.65/.011** | **-2.51/.016** |

### c. Characteristic path length (CPL)

Shortest path lengths were computed by defining edge distances as the inverse of weights[27, 28]:

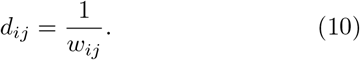

The characteristic path length was then defined as the average shortest path between all node pairs:

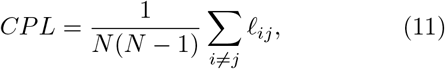

where *l*_*ij*_ denotes the length of the shortest weighted path between nodes *i* and *j*.

Our analysis showed, a subnetwork-dependent pattern for *CPL*. At the whole-brain level, MSE–Nets exhibited higher *CPL* than TS–Nets. In contrast, within the cortical and subcortical subnetworks, *CPL* was lower in MSE–Nets. Since *CPL* reflects integration (inversely)[2], these results suggest that MSE-based connectivity can yield reduced within-subnetwork distances (greater intra-system integration), while simultaneously increasing global distances at the whole-brain level. One possible explanation for this pattern is that MSE similarity between cortical and subcortical regions is comparatively weaker than intra-cortical or intra-subcortical similarity. In such a configuration, stronger within-subnetwork connectivity combined with relatively reduced cross-system similarity would naturally lead to shorter paths inside each subsystem but longer characteristic paths when the entire network is considered.

**TABLE III.** Statistical comparison of graph-theoretical measures between TS-based (TS-Nets) and MSE-based (MSE-Nets) functional brain networks within cortical across different age-ranges. Values are reported as *t*-statistics and corresponding FDR-corrected *p*-values (t/q) obtained from permutation-based *t*-tests. Results are presented for cortical and subcortical subnetworks in three age cohorts (21 −25, 26 −31, and 32 −37 years) across two independent dataset splits (Analysis 1 and Analysis 2). Graph measures include connection density (CD), clustering coefficient (CC), characteristic path length (CPL), Shannon entropy (S), and graph energy (H). Significant differences (*q <* 0.05) are indicated in **bold**.

| Age-range | | Density ($t/q$) | CC ($t/q$) | CPL ($t/q$) | H ($t/q$) | S ($t/q$) |
| --- | --- | --- | --- | --- | --- | --- |
| **TS-Cortex** |  |  |  |  |  |  |
| 21–25 | Analysis 1 | −0.59/.592 | −1.37/.409 | −1.31/.409 | 1.28/.409 | −0.63/.592 |
|  | Analysis 2 | −2.55/.097 | −1.90/.207 | −1.80/.207 | 2.35/.097 | −2.51/.097 |
| 26–31 | Analysis 1 | 1.17/.409 | −1.12/.409 | −1.12/.409 | 0.40/.684 | 0.89/.468 |
|  | Analysis 2 | 0.66/.624 | −1.22/.564 | −1.02/.564 | 0.25/.859 | 0.87/.581 |
| 32–37 | Analysis 1 | −0.96/.461 | −1.32/.409 | −1.79/.409 | 1.62/.409 | −1.19/.409 |
|  | Analysis 2 | −1.01/.564 | −0.47/.745 | −0.18/.859 | 0.69/.624 | −0.97/.564 |
| **MSE-Cortex** |  |  |  |  |  |  |
| 21–25 | Analysis 1 | 1.36/.209 | −0.74/.461 | −1.01/.374 | 0.93/.377 | −1.69/.123 |
|  | Analysis 2 | **2.93/.011** | −1.86/.071 | −1.67/.110 | 1.27/.201 | **−2.48/.021** |
| 26–31 | Analysis 1 | **5.88/ < .001** | **−4.03/.001** | **−4.58/ < .001** | **2.78/.009** | **−4.19/ < .001** |
|  | Analysis 2 | **4.31/.002** | **−3.28/.006** | **−3.69/.002** | **2.14/.038** | **−2.97/.011** |
| 32–37 | Analysis 1 | **4.23/ < .001** | **−2.44/.027** | **−2.91/.009** | 2.07/.061 | **−3.20/.006** |
|  | Analysis 2 | **2.99/.011** | **−2.61/.017** | **−2.80/.012** | **2.16/.038** | **−2.60/.020** |

**TABLE IV.** Statistical comparison of graph-theoretical measures between TS-based (TS-Nets) and MSE-based (MSE-Nets) functional brain networks within subcortical across different age-ranges. Values are reported as *t*-statistics and corresponding FDR-corrected *p*-values (t/q) obtained from permutation-based *t*-tests. Results are presented for cortical and subcortical subnetworks in three age cohorts (21 −25, 26 −31, and 32 −37 years) across two independent dataset splits (Analysis 1 and Analysis 2). Graph measures include connection density (CD), clustering coefficient (CC), characteristic path length (CPL), Shannon entropy (S), and graph energy (H). Significant differences (*q <* 0.05) are indicated in bold.

| Age-range | | Density ($t/q$) | CC ($t/q$) | CPL ($t/q$) | H ($t/q$) | S ($t/q$) |
| --- | --- | --- | --- | --- | --- | --- |
| **TS-Sub** |  |  |  |  |  |  |
| 21–25 | Analysis 1 | 0.63/.698 | 0.14/.896 | 0.33/.881 | −0.19/.896 | 0.61/.698 |
|  | Analysis 2 | −1.65/.317 | −1.61/.317 | −1.65/.317 | 1.54/.317 | −1.67/.317 |
| 26–31 | Analysis 1 | 2.04/.158 | 2.09/.158 | 1.94/.164 | −2.21/.158 | 2.07/.158 |
|  | Analysis 2 | 1.38/.317 | 0.57/.573 | 0.90/.401 | −1.00/.379 | 1.43/.317 |
| 32–37 | Analysis 1 | −0.98/.616 | −0.86/.616 | −0.85/.616 | 0.89/.616 | −1.08/.616 |
|  | Analysis 2 | −1.21/.317 | −1.16/.317 | −1.25/.317 | 1.17/.317 | −1.19/.317 |
| **MSE-Sub** |  |  |  |  |  |  |
| 21–25 | Analysis 1 | 0.31/.871 | −0.46/.871 | −0.20/.938 | 0.51/.837 | −0.04/.966 |
|  | Analysis 2 | **2.53/.029** | **−2.69/.012** | **−2.31/.029** | **2.90/.026** | **−2.51/.029** |
| 26–31 | Analysis 1 | 1.66/.371 | −1.75/.371 | −1.50/.385 | 2.20/.231 | −1.08/.591 |
|  | Analysis 2 | 0.35/.771 | −0.50/.731 | −0.49/.731 | 0.99/.450 | −0.11/.914 |
| 32–37 | Analysis 1 | 1.55/.374 | −0.98/.698 | −0.85/.724 | 0.60/.837 | −2.19/.231 |
|  | Analysis 2 | 2.10/.067 | −2.05/.069 | **−2.35/.029** | 1.47/.216 | −2.13/.067 |

### d. Graph energy (H)

*H* was computed from the eigenvalue spectrum of the weighted adjacency matrix[4, 29]:

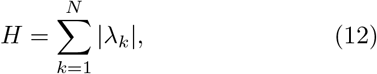

where *λ*_*k*_ are the eigenvalues of the adjacency matrix *A*. This measure reflects the overall spectral strength and is related to synchronization stability in the network.

According to our analysis, *H* also showed a network-dependent effect. TS–Nets exhibited higher energy than MSE–Nets at the whole-brain level and within the cortical subnetwork, whereas no significant difference was observed in the subcortical subnetwork. Interpreting energy as a spectral index related to global synchronization stability[4, 30], this pattern indicates that TS-Nets showed more stable synchronization in comparison to MSE-Nets (in the whole-brain and cortical levels).

### e. Shannon entropy (S)

Network-level *S* was defined over normalized edge weights[4, 31]:

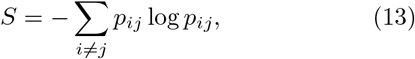

where

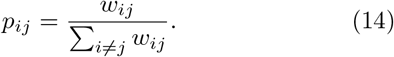

*S* quantifies the heterogeneity of the edge-weight distribution, reflecting the complexity of network connectivity[4, 30].

Our analysis showed that *S* patterns for both network types differed across subnetworks. MSE–Nets exhibited higher *S* than TS–Nets for the whole-brain network and within the cortical subnetwork, but lower *S* within the subcortical subnetwork. These findings suggest that MSE-based connectivity produces a more heterogeneous weight distribution globally and cortically, while the subcortical MSE-based connectivity becomes more concentrated (less heterogeneous) relative to TS-based connectivity.

## VI. SEX DIFFERENCES AND DISCRIMINATIVE SENSITIVITY OF NETWORK CONSTRUCTIONS

To evaluate the sensitivity of TS–Nets and MSE– Nets to biologically significant group differences, we examined sex-related effects in five graph measures (*CD, CC, CPL, H*, and *S*). As in the previous section, null models were used for normalization[16]. To improve the robustness of our results, we split the dataset into two independent halves and performed statistical analyses separately on each subset. Group comparisons were conducted using permutation t-tests[25], and all p-values were corrected using FDR.

### a. Whole-brain networks

At the whole-brain level, MSE–Nets exhibited robust and highly significant sex differences across all five measures in both Analysis 1 and Analysis 2. In contrast, TS–Nets showed largely non-significant differences, with only sporadic effects that failed to replicate consistently across both halves of the dataset (I; 3). Specifically, in the MSE-Nets, females exhibited higher *CD* and higher *H*, whereas males showed higher *CC*, higher *CPL*, and higher *S* (I; 3).

Higher *CD* in females indicates stronger average connectivity magnitude under the MSE-based similarity definition and higher *H* in females suggests greater overall spectral strength and enhanced global synchronization stability[30]. In contrast, higher *CC* and *CPL* in males reflects stronger local segregation and reduced global integration relative to females[2]. Finally, higher *S* in males implies a more heterogeneous and distributed edge-weight organization, reflecting greater complexity in connectivity pattern[30].

### b. Cortical network

Within the cortical subnetwork, MSE–Nets again demonstrated consistent and statistically significant sex differences across most measures in both analyses. Specifically, females exhibited a higher *CD* and higher *H*, while males showed a higher *CC*, a longer *CPL* and a higher *S* (I; 3). These results are consistent with whole-brain network results.

Within the cortical subnetwork, TS–Nets also revealed significant sex differences for selected measures. Specifically, *CC, CPL*, and *H* were significant in both independent analyses, indicating reproducible sex-related effects in cortical segregation and integration[2] under the timeseries–based connectivity definition (I; 3). These effects were limited to a subset of measures and did not extend consistently to *CD*, and *S*.

### c. Subcortical network

In the subcortical subnetwork, MSE–Nets again revealed robust sex differences for most measures. Males showed higher *CD* and higher *H*, while females exhibited higher *CC* and *S*. In contrast to the cortical and whole-brain networks, *CPL* in the subcortical network were not consistently significant across analyses.

These findings indicate that, at the subcortical level, males exhibit stronger overall coupling strength and spectral stability, whereas females demonstrate greater local segregation and higher connectivity heterogeneity. The absence of stable *CPL* differences suggests that integration-related effects[2] are less pronounced within the subcortical system compared to cortical or wholebrain organization.

### d. Age-stratified analyses

To further evaluate the robustness of our findings, we repeated the analyses across three distinct age cohorts: 21–25, 26–31, and 32–37 years. To maintain brevity, we report only the results for whole-brain network features. detailed results for cortical and subcortical network features across these age ranges are provided in the III, and IV.

Consistent with previous whole-sample findings, TS– Whole networks showed no significant sex effects within any age bin. In contrast, MSE–Whole networks exhibited multiple significant differences across age ranges, particularly in the 26–31 and 32–37 groups, with several effects replicating across both dataset splits (II; 3.

## VII. DISCUSSION

The present study introduces a multiscale framework for functional brain network construction in which interregional similarity is defined through correlations between entropy profiles rather than direct temporal synchrony. Our findings demonstrate that this formulation leads to systematic differences in network topology, mesoscale organization, and spectral properties compared to conventional time-series–based networks.

MSE-based networks consistently exhibited stronger modular organization and enhanced local segregation, while simultaneously reshaping global integration patterns. This suggests that similarity in multiscale dynamics captures aspects of functional organization that are not accessible through direct synchrony alone[9, 10]. Importantly, the observed subnetwork-dependent effects indicate that multiscale similarity differentially constrains intra- and inter-system interactions, particularly between cortical and subcortical regions.

A central result of this work is the substantially higher sensitivity of MSE-based networks to biologically meaningful variability. Sex-related differences were robust, consistent across independent dataset splits, and reproducible across age ranges in MSE-based networks, whereas TS-based networks showed limited and inconsistent effects. This indicates that multiscale representations provide a more informative basis for detecting structured variability in brain organization[9, 10].

## VIII. CONCLUSION

In summary, defining functional connectivity through MSE similarity provides a principled alternative to conventional synchrony-based approaches. By embedding scale-dependent and nonlinear properties of neural dynamics into network construction, this framework reveals organizational features and group differences that remain largely undetected in standard formulations. These results highlight the importance of multiscale representations in network neuroscience and suggest that MSE-based connectivity may offer a more sensitive and biologically meaningful characterization of brain function.

## IX. CODE AND DATA AVAILABILITY

The neuroimaging data used in this study were obtained from the Human Connectome Project (HCP) (Young Adult S1200 release), available at https://www.humanconnectome.org/. Use of the HCP data is subject to the acceptance of the HCP Data Use Terms.

Custom MATLAB scripts for calculating graph energy, entropy, and permutation-based statistical analyses are available at the following GitHub repository: https://github.com/AHGhaderi. For the calculation of other graph-theoretical metrics, including modularity, CC, and CPL, we utilized the Brain Connectivity Toolbox (BCT) (https://brain-connectivity-toolbox.net/). The MSE algorithm was implemented based on the framework proposed by [20] available at https://www.mathworks.com/matlabcentral/fileexchange/62706-multiscale-sample-entropy. All processed connectivity matrices and analysis pipelines used to generate the results in this study are available upon request from the corresponding author.

